# Environmentally Relevant Polylactic Acid Microplastics from 3D Printing Induce Germline Apoptosis and Reproductive Decline in *Caenorhabditis elegans*

**DOI:** 10.64898/2026.07.24.740412

**Authors:** Chiara Angelyn O. Maldonado, Dana Farley, David M. Mares, Ryan J. Rutledge, Maria F. Gamez, Laila R. Sosa, Viviana Palacios, Amy Gutierrez, Ryan L. Peterson, Jennifer C. Harr

## Abstract

Bio-based plastics, such as polylactic acid, offer an alternative to petroleum-based plastics and a prospect to address the plastic pollution problem. While mounting evidence suggests that microplastics pose a human health threat, the risk posed by bio-based plastics remains unknown. Here, we use *Caenorhabditis elegans* to investigate the effects of secondary microplastics from 3D-printed polylactic acid on fertility and lifespan. We have created and characterized microplastics from a 3D-printed item using cryogenic milling. Using these microplastics, we exposed *C. elegans* and assessed lifespan, reproduction, and various stress responses. Our studies demonstrate that exposure to 1 µg/L polylactic acid microplastics reduces fertility and alters the gonad structure. When examining germline integrity, we find chromosomal disorganization in the gonad after polylactic acid microplastic exposure and increased apoptotic cell death, which correlates with a DAF-16/FOXO and gst-4 oxidative stress responses. While lifespan has been observed to decrease with exposure to microplastics of different polymer types, we did not observe a change in lifespan with exposure to polylactic acid microplastics. The germline is likely more sensitive than somatic tissues under our exposure conditions. The reduction in fertility is driven by alterations to the germline, characterized by chromosome aberrations, oxidative stress, and cell death.

## 1. Introduction

While bio-based plastics offer an alternative to petroleum-based plastics and help address the pervasive problem of plastic pollution, their health risks remain understudied. Bio-based plastics hold promise for reducing plastic waste due to their ability to degrade and their perceived environmental and health benefits over conventional plastics [1, 2].

The first bio-based plastic was synthesized in the 1930s, though its widespread use has only recently increased significantly [1]. Lactic acid (LA) was first discovered by the chemist Wilhelm Scheele and was later polymerized by Wallace Hume Carothers as polylactic acid (PLA) [1]. Its early production was cost-prohibitive, and it did not gain widespread use until the 1960s, as absorbable sutures and drug-delivery matrices in biomedicine [3-5]. Since being approved by the FDA for biomedical uses in the 1970s, PLA and other bio-based plastics have been used in food packaging [2, 6, 7] and, more recently, as 3D-printing filaments [8]. PLA currently accounts for the largest share of filaments used in the 3D printing landscape, and its use is expected to continue to increase [8].

PLA is a bio-based, biodegradable plastic derived from fermented plant sugars [1]. It has advantages such as reduced use of conventional petroleum-based plastics and its use in biomedical and food applications. The biodegradation of PLA occurs through hydrolysis of its ester backbone [9]. Under high temperatures and humidity, this process produces lactic acid oligomers and finally, the lactic acid monomer, which can be metabolized by some microbial organisms into carbon dioxide, water, and biomass [9, 10]. However, it does not fully degrade under most real-world environmental conditions and can lead to the production of PLA microplastics [10]. As PLA requires very specific conditions to degrade (i.e., humidity (> 50%) and temperature (> 55 °C) [9], it can persist in the environment [11] and produce secondary microplastics (MP; particles < 5 mm [12]) and nanoplastics (NP; particles < 1 µm [12]). MPs and NPs of a wide variety of polymer types have been found in animal and human tissues [13-16]. There is overwhelming evidence for the detrimental effect of MPs on health and development. In a wide range of animal model systems, MPs and NPs lead to reduced lifespan and reproductive decline and can act as carriers to enhance their detrimental effects [17-20].

Much of the evidence on MP and NP exposure relies on pristine primary microplastic stocks because they were readily available [21, 22]. However, these pristine MPs do not adequately represent the secondary microplastics found in the environment. Evidence shows that MPs with irregular size, shape, and surface texture have more detrimental effects than their pristine counterparts [21, 23, 24]. Irregular secondary MPs are created by the breakdown of larger items, wear from products during use, or generation in waste management [25]. This breakdown can occur via UV exposure, mechanical wear, or the fragmentation of larger plastic particles [25]. Additionally, primary and secondary MPs can contain additives from the manufacturing process or absorb or carry chemicals they may pick up from the environment, which can be transferred to animals after ingestion [17, 26]. Thus, the study of secondary (irregular) MPs has direct implications for human health and remains an understudied area of MP and NP exposure research compared with pristine plastic counterparts.

The assumption that PLA is “safe” is being challenged with emerging evidence that PLA-MPs can cause neurotoxicity, intestinal damage, inflammation [27, 28], and reduce fertility [29-31]. Here we use the well-established *Caenorhabditis elegans* (*C. elegans*) model [32] to investigate the effects of irregular secondary PLA-MPs on reproduction and lifespan. We made PLA-MPs from an item produced with a 3D printer to mimic common types of waste that could lead to the production of MPs through improper disposal. We chose to produce our secondary MPs in-house using a mechanical method that does not introduce additional chemicals. This allowed us to examine the effects of heterogeneous shape, surface texture, surface structure, and size. We chose PLA-MP exposure levels for this study (1 µg/L) that are within the range of other studies asking about MP effects, and that mimic concentrations determined from human samples, such as blood [13, 19, 29, 33, 34]. We hypothesized that a heterogeneous, irregular pool of PLA-MPs would lead to physiological defects in *C. elegans*. We find that exposure to PLA-MPs reduced brood size, altered gonad development, induced apoptotic cell death, and led to oxidative stress response in the germline.

### 2. Materials and Methods

### 2.1. Creation of Secondary Microplastics

A plastic cone was printed on an A1 Mini 3D printer (Bambu Labs). Orange 1.75 mm PLA filament (PrintBed, U.S. NatureWorks, Plymouth, MN, USA) was used to make the item. A 2.5 g plastic cone fragment was collected and placed in a Freezer/Mill CG-200 (Cole-Parmer, Metuchen, NJ, USA). Using the thermoplastic setting, the following protocol was used: 15-minute precool, 12 grinding cycles (2 minutes each), 2 minutes cool, and a rate of 12 cycles per second. This cycle was repeated 3 times. The plastic particles were then placed in a sieve stack (starting with the largest, Sieve No. 40 [425 µm], Sieve No. 60 [250 µm], Sieve No. 80 [180 µm], Sieve No. 100 [150 µm], and ending with the smallest, Sieve No. 63µm [63 µm], Standard Sieves Dual Manufacturing stainless steel mesh, ASTM E11 Compliant Sieves, Franklin Park, IL, USA), and the stack was manually shaken for 10 minutes. Fragments of MPs that passed through the 63-micron opening were collected for use in experiments.

### 2.2. Characterization of Microplastics

ATR-IR analysis was used to determine the plastic and chemical profile. Using the Bruker Alpha Platinum ATR, a scoop of PLA-MPs was placed on the crystal part of the instrument and scanned at a resolution of 4 cm-1, 16 scans, and a range of 400-4000 cm-1. MP size and surface were characterized by Scanning Electron Microscopy (SEM; JEOL JSM-6010 PLUS/LA, Peabody, MA, USA). Samples were sputter-coated with 2 nm gold particles prior to imaging (Quorum Technologies EMS150T ES, Lewes, UK). MP size (area, smallest width, longest width, and thickness) was measured using ImageJ software.

### 2.3. Preparation of E. coli OP50 food and PLA Microplastics

MPs were suspended in *Escherichia coli* (*E. coli*, strain OP50, grown in LB broth), seeded on Nematode Growth Media (NGM) plates (Teknova, N1105, Hollister, CA, USA), and allowed to dry overnight before adding *C. elegans* to the plates for exposure. To prepare PLA-MP *E. coli* suspensions, PLA MPs were suspended in a 1% solution with 1x phosphate-buffered saline (PBS), then diluted into *E. coli* OP50. At each dilution step, the solution was sonicated for 15 min (Emerson Branson 3800, Danbury, CT, USA). Wide-bore tips were used to transfer liquid. In-house prepared PLA-MPs were used for all experiments.

### 2.4. Maintenance of C. elegans

*Caenorhabditis elegans (C. elegans)* strains, N2 (wild type), TJ356 (zIs356 [daf-16p::daf-16a/b::GFP + rol-6(su1006)]), CL2166 (dvIs19 [gst-4p::GFP::NLS] III), and MD701 (bcIs39 [lim-7p::ced-1::GFP + lin-15(+)] that were used in this study were provided by the Caenorhabditis Genome Center. *C. elegans* were grown on NGM plates seeded with *E. coli* at 20 °C.

### 2.5. Exposure Design

Bleach-synchronized L1 larvae were exposed to either *E. coli* or *E. coli* containing secondary 1 µg/L PLA-MPs in *E. coli* on NGM plates made within 48 hours. At least 2-3 biological replicates were performed for each assay. For stress assays and gonad chromosome analysis, L1-synchronized larvae were placed onto each condition, incubated for 72 hours at 20 °C, and imaged. For the gst-4 GFP oxidative stress positive control, *C. elegans* were exposed to either M9 or 0.1 mM paraquat in M9 and incubated at 20°C for 1 hour before dissecting the gonads and imaging live.

### 2.6. Lifespan Assay

Lifespan assays were performed at 20 °C. Approximately 50 synchronized L1 larvae were placed on the corresponding treatment conditions for 48 hours and then transferred to new plates containing the same treatment with 100 µM 5′-fluorodeoxyuridine (RPI, 50-91-9, Mount Prospect, IL, USA). *C. elegans* were counted as dead or alive every day until the number of live *C. elegans* reached zero.

### 2.7. Evaluation of Reproductive Toxicity and Incidence of Males

To determine the total number of eggs laid per *C. elegans* and the percentage hatched, bleach-synchronized L1 larvae were seeded onto either control (*E. coli*) or 1 µg/L PLA-MPs in *E. coli* OP50. 48 hours after seeding, L4 larval *C. elegans* were then singled out onto respective plates. The following day, L4 singling is repeated from the previous day’s plate, and both plates are kept. After 24 hours, L1s and embryos are counted from the first day of singling, then repeated every day, counting the L4-free plate, until egg laying stops (around 6-7 days). Plates are kept at 20 °C for an additional 24 hours, until *C. elegans* are L4s/young adults, and the total number of males per plate is counted.

### 2.8. Imaging and Microscopy

All fluorescent and bright-field images were captured using an EVOS M5000 (Invitrogen, Bothell, WA, USA). Specifically, Carnoy’s-fixed and Hoechst-stained whole *C. elegans*, images were taken with a Zeiss LSM 910 (Zeiss, Jena, Germany). For live imaging of the whole *C. elegans (DAF-16::GFP and CEC-1::GFP)* were transferred to an agarose pad on a slide and immobilized with levamisole (Vector Laboratories, SP-5000-18, Newark, CA, USA). For live imaging of dissected gonads (gst-4p-GFP expression), gonads were dissected from *C. elegans* and imaged. All image processing and analysis were performed using ImageJ software (version 1.54). For the gonad area and length, FIJI (version 2.9.0) was used for analysis.

### 2.9. Activation of Stress Response

To determine whether PLA-MP exposure induces stress activation, the strain TJ356, expressing DAF-16 fused to GFP, was exposed to PLA-MPs as described. Live imaging was completed within 5 minutes of paralysis. Each *C. elegans* was classified by the localization of DAF-16::GFP expression (cytoplasmic, intermediate, or nuclear).

### 2.10. Oxidative Stress Assay

To determine whether PLA-MP exposure leads to oxidative stress, the strain CL2166, expressing GFP under the gst-4 promoter, was exposed as described. This reporter strain has GFP fused to the promoter of glutathione S-transferase 4, and oxidative stress induces GFP expression. Following exposure, *C. elegans* were transferred into a drop of M9 on a positively charged slide. The *C. elegans*’ gonads were dissected and isolated from their carcasses to minimize background autofluorescence. Dissected gonads were imaged within 10 minutes of dissection. Imaging settings were consistent across samples. Fluorescent intensities were determined in ImageJ software. A set of gst-4p::GFP *C. elegans* was treated for 1 hour with or without 50 mM paraquat (PQ) (in M9) and was used as positive and negative controls, respectively.

### 2.11. Germ Cell Apoptosis

To determine if PLA-MP exposure leads to cell death and crypt cell formation in *C. elegans*, wild-type *C. elegans* were exposed to 1 µg/L PLA-MPs in *E. coli* or *E. coli* only for 72 hours. Adult *C. elegans* were washed from plates and incubated with 25 μg/mL of acridine orange (Abcam, ab270791, Waltham, MA, USA). Tubes were wrapped in foil to protect from light and placed on a rotator at room temperature for 1 hour. *C. elegans* were then transferred to NGM plates with *E. coli* and protected from light for 1 hour. Stained *C. elegans* were then mounted on 4% agarose pads with levamisole and imaged. MD701 *C. elegans* (expressing a CED-1::GFP fusion protein in sheath cells) were exposed to 1 µg/L PLA-MPs in *E. coli* or *E. coli* only. L1-synchronized larvae were placed onto each condition, and live-imaged 72 hours later.

### 2.12. Chromosome organization

After PLA-MP and control exposure (as described above), adult *C. elegans* were picked and placed onto a concave slide containing a drop of M9. Bacteria were washed away by pipetting off and discarding the original M9 drop, then replacing it with the new M9, ensuring the *C. elegans* were untouched. The *C. elegans* were then pipetted onto a positively charged slide with a drop of M9 + 1 mM levamisole. Within 2 minutes, the gonads were dissected using a 27G needle to cut off the heads, just below the pharynx. The excess paralytic liquid was pipetted out and wicked away. Following dissection, gonads were fixed in 4% paraformaldehyde (PFA) in 1x phosphate-buffered saline (PBS) for 20 minutes. After incubation, excess liquid was removed. The sample was then washed three times for 3 minutes each with PTX solution (10x PBS, 10% Triton X). Next, *C. elegans* were stained with Hoechst for 5 minutes, followed by one wash with PTX. Mounting media was added prior to adding a coverslip. These were then imaged using an EVOS fluorescence microscope and subsequently quantified.

### 2.13. Gonad morphology

Analysis of adult gonad morphology after PLA-MP and control exposure (as described above) was performed by fixing adult *C. elegans* with Carnoy’s fixative (6 ethanol: 3 chloroform: 1 glacial acetic acid). *C. elegans* were wicked into a minimal volume of M9 buffer on a microscope slide, and excess liquid was wicked away. *C. elegans* were then fixed by dropping 2-3 drops of Carnoy's fixative onto the slide from a height of 2 inches, and the slide was allowed to dry. *C. elegans* were then rehydrated in 20 μl of M9 for 1 hour. After wicking away excess M9, 10 μl of staining solution (1:10,000 Hoechst 33342 (20 mM, Thermo Scientific 62249)) was added and incubated for 5 minutes. Excess staining solution was wicked away, and the remaining area was washed once with M9. Mounting medium was added, and the slide was covered with a coverslip. Slides were left overnight at 4 °C before imaging. Carnoy’s-fixed and Hoechst-stained *C. elegans* were used to measure gonad area and length.

### 2.14. Statistical Analysis

Statistical analysis and figure design were performed with GraphPad Prism 8 (GraphPad Software version 11, San Diego, CA, USA). Data were checked for normality between treatments, and statistical analysis was performed using a Mann-Whitney test (brood size, oxidative stress, apoptosis, and gonad measurements). A chi-square test was performed to evaluate the localization of DAF-16 in TJ356 using raw frequency counts, with data presented as percentages. Fisher’s exact test with 95% C.I. was performed to analyze the raw counts of Hoechst staining data. Experiments were all carried out with a minimum of 2-3 biological replicates (N) and the number of total *C. elegans* (n) examined, which were used in statistical analysis as described.

## 3. Results and Discussion

### 3.1. Creation and Characterization of PLA Secondary Microplastics

Secondary polylactic acid (PLA) microplastics (MPs) were produced from discarded 3D-printed items by cryo-milling and size-selected using ASTM E11-compliant standard sieves to 63 µm or smaller (Figure 1a). To determine the particle size and surface characteristics, scanning electron microscopy (SEM) was used to image PLA-MPs. PLA-MPs were irregular in size and shape, with an overabundance of flat irregular shapes (Figure 1b). Quantification of particle size using the shortest and longest surface (Figure 1c, Supplemental Figure 1a) showed a variety of sizes ranging from 1-150 µm (Figure 1c, Supplemental Figure 1). The area of the largest surface visible showed a broad distribution, with a large portion of the population skewing towards a smaller overall area (Figure 1d). In our work, we chose to generate environmentally realistic secondary MPs from plastic waste, rather than relying on homogeneous commercially available MPs. Secondary MPs can be produced in the laboratory with a variety of techniques, such as Ultraviolet (UV) exposure, chemical treatment, and mechanical breakdown [23, 35-37]. Mechanical methods are a top-down approach that use physical fragmentation of plastic through processes such as cutting, grinding, milling, and cryogenic treatment. The particles produced by these processes mimic the irregular characteristics of environmental plastics [35, 37]. The fragments we produced had irregular and heterogeneous morphologies, as observed with mechanical breakdown of other polymer types [37], and the integration of sieves allows us to narrow the range of particle sizes tested [37, 38]. It is important to note that during our suspension of MPs in the *C. elegans* food source, we perform multiple sonication steps. Sonication has been used to fragment materials and create MPs [35]. Therefore, in our final suspension of PLA-MPs presented to *C. elegans*, there is likely a larger proportion of smaller particles than we determined by SEM, as this step was performed prior to sonication.

**Figure 1.**
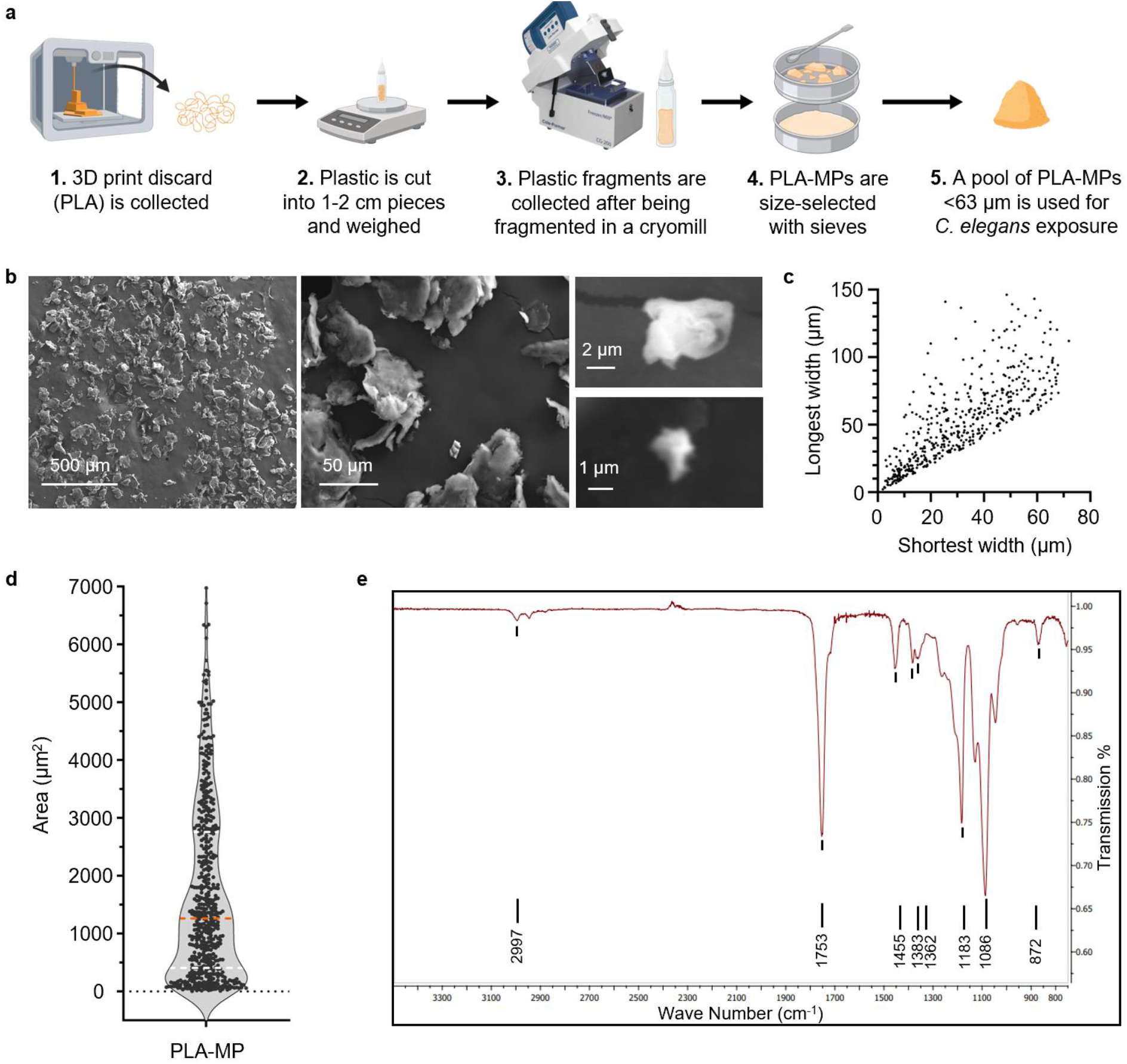
Creation and characterization of secondary MPs from 3D printer PLA waste. (**a**) Diagram of the process of making a less than 63 µm pool of PLA-MPs. Created with BioRender.com. (**b**) Representative SEM images of PLA-MPs to determine size, shape, and surface characteristics. (**c**) PLA-MP particle size distribution, n = 510. (**d**) PLA-MP particle area distribution, n = 510. (**e**) ATR spectrum of PLA-MP.

**Figure 2.**
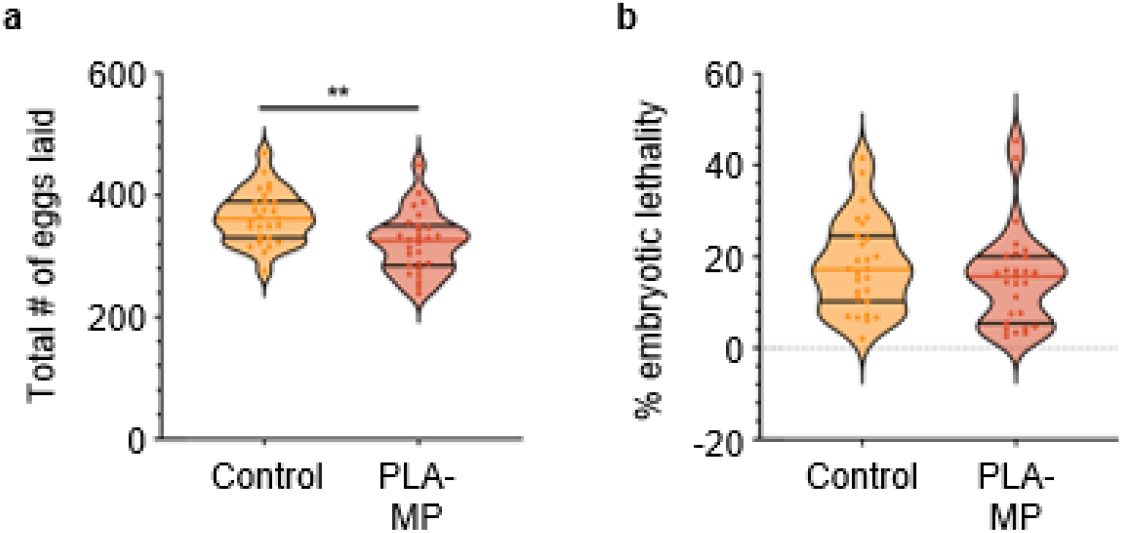
Total eggs laid and embryonic lethality of *C. elegans* exposed to PLA-MPs. (**a**) Number of eggs laid per *C. elegans* with *E. coli* only or with PLA-MPs. N = 4, n = 27, 27. (**b**) Percent embryonic lethality. Violin plots show the mean with the standard deviation. Mann-Whitney (two-sample) tests were used for statistical analysis (** *p* ≤ 0.01).

MP exposure can be through ingestion, dermal contact, and inhalation [39, 40]. MPs of various polymers and mixtures have been detected in prepared food products, raw fruits and vegetables, drinking water, and beverages [41-44]. Many studies of MP exposure focus on ingestion. When considering ingestion as a source of exposure, *C. elegans* is limited by the size of its mouth (∼4 µm in adults) [45, 46]. The provided food source, *E. coli*, is approximately 1 µm in size. In the wild, *C. elegans* will consume a variety of microbial food sources, though they show a preference for smaller food particles, and ingestion of larger items can lead to reduced feeding or obstruction [45]. Regarding MPs, *C. elegans* have been shown to ingest those up to 5 µM in diameter [47]. Analysis of the size distribution of the produced PLA-MPs indicated that approximately 3% of the particles are small enough to be ingested (≤ 5 µm, the smallest particle size). At an exposure of 1 µg/L, the relative ingestion exposure is approximated to be 30 ng/L. A limitation of this estimation is a potential bias towards imagining larger particles, as they are more easily identified. Ingestion of an irregular pool of PLA-MPs may lead to ingestion of larger particles (> 5 µm). Thus, causing inefficient ingestion and indirect toxicity through feeding interference or dermal exposure.

Next, we examined the chemical properties of the 3D-printer-extruded and cryo-milled PLA-MP particles. ATR-FT-IR spectra showed a similar IR profile (Figure 1e) to that observed for reported PLA samples [6]. Some plastic polymers undergo physical changes that alter their ATR/FTIR spectra after UV degradation, a process that facilitates the secondary creation of MPs [23, 48]. We did not observe any chemical differences, which is consistent with studies showing that 3D-printed PLA materials are not altered by the extraction and heating processes used in 3D printing [49]. We chose to create PLA-MPs using a cryogenic mill to mimic mechanical degradation without chemical or UV treatment. This allowed us to examine the consequences of a non-pristine microplastic particle without adding a chemical treatment or additive.

### 3.2. C. elegans Exposed to PLA-MPs Leads to Reduced Fertility

To determine whether PLA-MPs affect reproductive toxicity, L1-stage *C. elegans* were continuously exposed to 1 µg/L PLA-MPs. There was a significant decrease in the total number of eggs laid when exposed to PLA-MPs (*p = 0.0021*) (Figure 3a). However, there was no significant change in embryonic lethality with PLA-MP exposure (Figure 3b, Supplemental Figure 2a, b). MPs or various polymers at similar concentrations have been shown to reduce brood size [17, 18, 23, 29, 50]. In contrast to our results with a pool of irregular PLA-MPs ranging from 1-150 µm in diameter, no reduction in brood size was seen with 1 µg/L exposure to ∼3 µm pristine PLA-MPs, whereas higher concentrations (10 and 100 µg/L PLA-MPs) reduced brood size [29]. While we did not observe a change in embryonic lethality, as was observed with pristine PLA-MPs at higher concentrations, we saw a marked decrease in brood size at a significantly lower exposure level. This enhanced effect is likely due to morphological differences between irregular secondary MPs and pristine MPs. Multiple studies show that irregular MPs have a more detrimental effect on fertility than their pristine counterparts. Shao et al. demonstrated that secondary MPs produced via UV irradiation reduced brood size compared with pristine PLA-MPs [24]. Compared with our cryo-milled PLA-MPs of broad size and no chemical modifications (as observed by ATR), this work showed that the uniform-sized PLA-MPs produced by UV irradiation were slightly smaller, had a more rugged surface, and exhibited altered chemical bonds when compared to those not treated with UV [24]. While there are differences in the production of secondary MPs (UV irradiation vs. mechanical) and distribution by particle size (more homogeneous vs. heterogeneous), we note that the pool of ingestible PLA-MPs in our study was significantly smaller than that used in Shai et al.’s study. Therefore, the highly irregular shape of cryo-milled MPs may be leading to reduced fertility.

**Figure 3.**
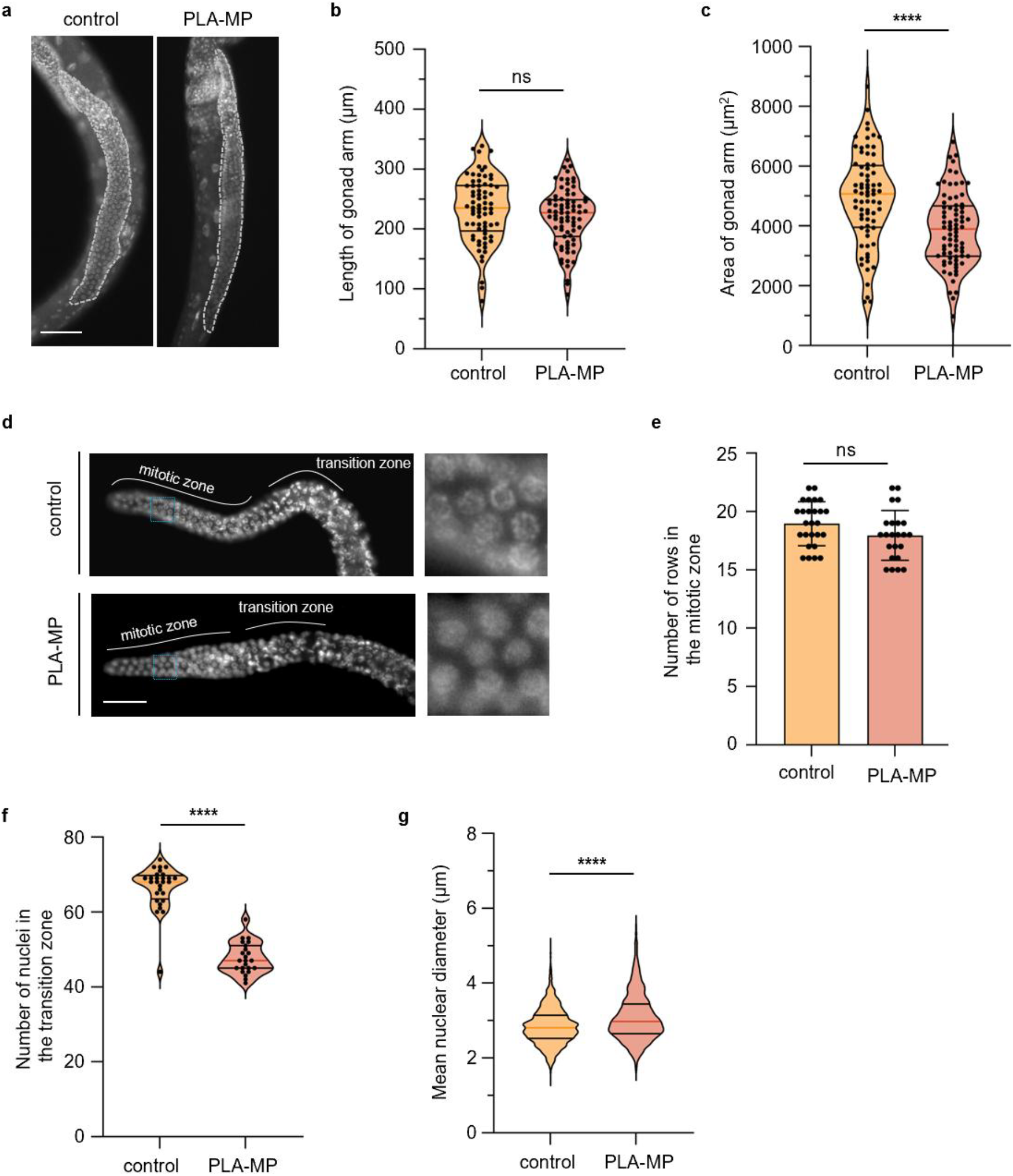
Alteration of gonad development with exposure to PLA-MPs. (**a**) Representative images of Hoechst staining in whole adult *C. elegans* fed *E. coli* without (control) and with PLA-MPs. The dotted line indicates the measured region of the gonad. Scale bar = 50 µm. (**b**) Quantification of the length of the gonad arm N = 3, n = 70, 75. ns = not significant. (**c**) Quantification of the area of the gonad arm N = 3, n = 72, 76. (**d**) Representative images of Hoechst staining in dissected gonads from adult *C. elegans* fed *E. coli* without (control) and with PLA-MPs, left whole gonad, right zoom in on pre-meiotic nuclei. Scale bar = 25 µm. (**e**) Quantification of the number of rows in the pre-meiotic zone. N = 2, n = 28, 20. (**f**) Quantification of the number of nuclei in the transition zone. N = 2, n = 28, 20. (**g**) Quantification of the mean nuclear diameter (μm) in *C. elegans* following control or PLA-MP exposure. N = 3, n = 1949, 1992 (8-10 gonads per condition. **** *p* < 0.0001 by the two-tailed Mann-Whitney test, 95% C.I.

### 3.2. PLA-MP Exposure Alters Gonad Development

Given the marked decrease in reproductive capacity, we next asked whether there were alterations in germline development in *C. elegans* exposed to PLA-MPs that might explain the reduced fertility. Exposure of *C. elegans* to 1 µg/L PLA-MPs led to a reduced gonad area (*p* < 0.0001) and a non-significant reduction in the length of the gonad (Figure 3a, b, c). Additionally, PLA-MP exposure did not alter the number of rows of mitotic cells per gonad, but it did reduce the number of cells in the transition zone (*p* < 0.0001; Figure 3d, e, f). PLA-MP exposure also increased the mean nuclear diameter of pre-meiotic nuclei (*p* < 0.0001; Figure 3d,f). Shao et al. showed similar reductions in gonad development with exposure to higher levels (10 and 100 µg/L) of uniform 3 µm-diameter PLA-MPs [29]. 1 µg/L 3 µm-diameter PLA-MPs did not affect gonad development in their study [29]. These differences are likely due to the differences in MP size, shape, and surface characteristics. Another study by Shao et al., comparing pristine and UV-aged PLA-MPs, demonstrated greater toxicity to reproductive capacity and gonad development in UV-aged PLA-MPs, which exhibited altered chemical and physical properties [24]. While we did not observe changes in the chemical properties of our cryo-milled PLA-MPs, they exhibit a rough surface structure and irregular shapes (Figure 1), as seen in UV-aged MPs, suggesting that many of the defects may be due to alterations in surface structure and the irregular size of secondary MPs. Enhanced reproductive defects and alterations in gonad development with exposure to irregular MPs, compared with pristine MPs, are consistent with results observed in studies using weathered PLA-MPs and other types of MPs [23, 30, 51]. The observed decrease in gonad size (Figure 3) and germline apoptosis (Figure 5) parallels those observed with polystyrene (PS) MPs [52]. Furthermore, across multiple species, MP toxicity leads to altered gonad structure, development, and reproductive capacity [18], as seen in other non-MP-related toxicity studies [53, 54].

### 3.3. PLA-MP Exposure Leads to an Increase in Chromosome Organization Defects and Apoptosis in the Gonad

Reduced fertility in *C. elegans*, leading to gonad defects, can be attributed to defects in chromosome organization and apoptosis in the gonad [55, 56]. Fertility defects can result from errors in meiosis leading to autosomal chromosome segregation and the formation of aneuploid gametes in *C. elegans* [57]. To assess whether defects in chromosome organization may lead to reduced oocyte development and contribute to reduced fertility, we examined the gonads of PLA-MP-exposed *C. elegans* stained with Hoechst. Three categories were defined to represent chromosomal defects that can serve as indicators of DNA damage [55]. Laggers are categorized as leptotene/zygotene-like nuclei commonly found in the transition zone, yet they persist into the pachytene stage [55]. While aggregates are multi-nuclei clumps and gaps are areas of reduced nuclear density [55]. With exposure to PLA-MPs, we observe a significant increase in aggregates (*p* = 0.0039) and laggers (*p* = 0.041) (Figure 4). However, there was no significant increase in the number of gaps (Figure 4). When examining *C. elegans* with more than one type of defect, 31.4% of the gonads observed had multiple defects after PLA-MP exposure, as compared to 7.9% in the control (*p* = 0.0042). Therefore, significantly more *C. elegans* with multiple defects were observed after PLA-MP exposure.

**Figure 4.**
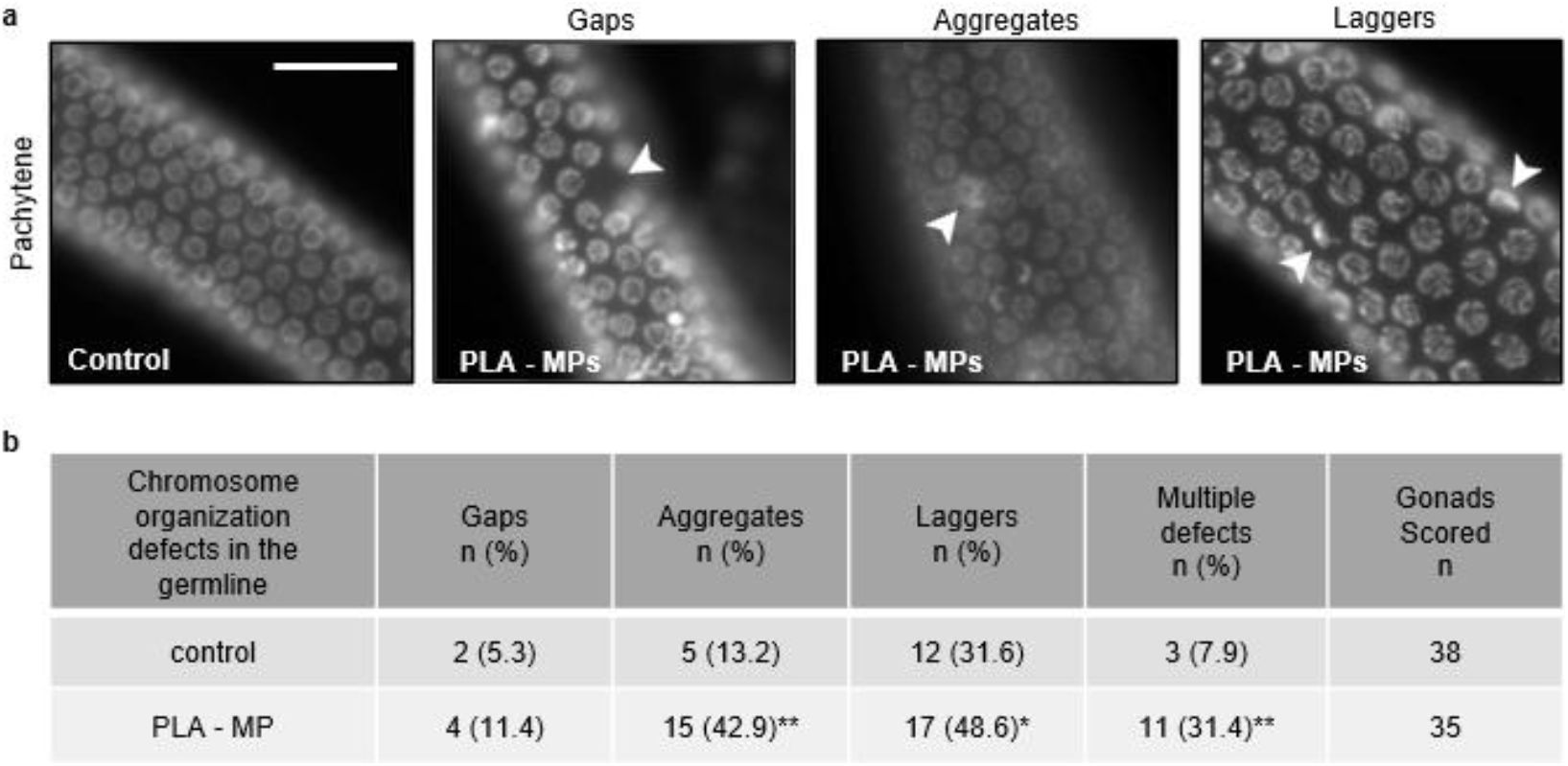
Observed defects in the chromosomal organization of *C. elegans* germline. (**a**) Representative images of Hoechst staining in adult *C. elegans* fed *E. coli* without (control) and with PLA-MPs. Scale bar = 25 µm. (**b**) Quantification of defects observed (gaps, aggregates, laggers) in Hoechst-stained gonad nuclei in *C. elegans* with exposure to their regular *E. coli* diet only or with 1 µg/L PLA-MPs. N = 3, n = 38, 35. Fisher’s exact test was used for statistical analysis (* *p* ≤ 0.05, ** *p* ≤ 0.01).

Shin et al. and Henderson et al. report similar germline chromosomal abnormalities resulting from exposure to soluble chemicals (phthalates and other toxicants) [55, 56]. Additionally, they observed chromosome non-disjunction due to some chemical exposures [55, 56]. In *C. elegans*, nondisjunction of the X chromosome during meiosis leads to the formation of males [58]. We measured the incidence of males in the offspring of *C. elegans* exposed to 1 µg/L PLA-MP and observed no increase compared with the control (Supplemental Figure 2c, d). Nonetheless, we observe defects in meiotic chromosome organization that may contribute to reduced fertility with PLA-MP exposure. Our results on defects in chromosome organization in the meiotic germline of PLA-MP-exposed *C. elegans* are consistent with evidence that MPs are genotoxic across species. Polystyrene microplastics (PS-MPs) delay porcine oocyte maturation by disrupting spindle architecture and metaphase I chromosome alignment [59]. PS-MPs similarly led to chromosomal instability in mammalian cells by increasing centriole numbers and altering microtubule-kinetochore attachments [60]. Specifically in *C. elegans*, PS-MP exposure leads to oocyte-specific chromosome aberrations and germline apoptosis [61]. Together with established germline toxicity assays showing chemically induced meiotic chromosome defects in *C. elegans*, these findings support the idea that secondary PLA-MPs from 3D-printed materials are genotoxic in the germline, perturbing chromosome organization during meiosis and contributing to reproductive decline.

Chromatin that has collapsed in the late pachytene region has previously been found to correlate with apoptosis in germ cells. Moreover, chromosome disorganization has been linked to DNA damage from exposure to soluble toxicants [55, 56], and MP studies observe DNA-damage-induced germline apoptosis [29, 51, 61]. Therefore, we next examined the germline in PLA-MP-exposed *C. elegans* for cell death and apoptosis. To determine whether PLA-MP exposure increases germ cell apoptosis, we observed germ cell corpses (apoptotic bodies) using acridine orange staining and monitored the expression of a functional CED-1::GFP fusion protein in the sheath cells of a reporter *C. elegans* strain. There was a significant increase in acridine orange-stained apoptotic cells when exposed to PLA-MPs compared to the control (*p*< 0.0001) (Figure 5 a, b). In addition, there is an increase in cell corpses when exposed to PLA-MPs (*p = 0.0042*) (Figure 6c, d). Pristine uniform 3 µm PLA-MPs (10 and 100 µg/L) led to germline apoptosis, while 1 µg/L PLA-MP did not [29]. Consistent with our results, UV-aged PLA-MPs induced germline apoptosis across all exposure levels (1, 10, and 100 µg/L) [30]. This indicates that irregular secondary PLA-MPs cause detrimental germline defects and apoptosis at lower exposure levels than pristine PLA-MPs do.

**Figure 5.**
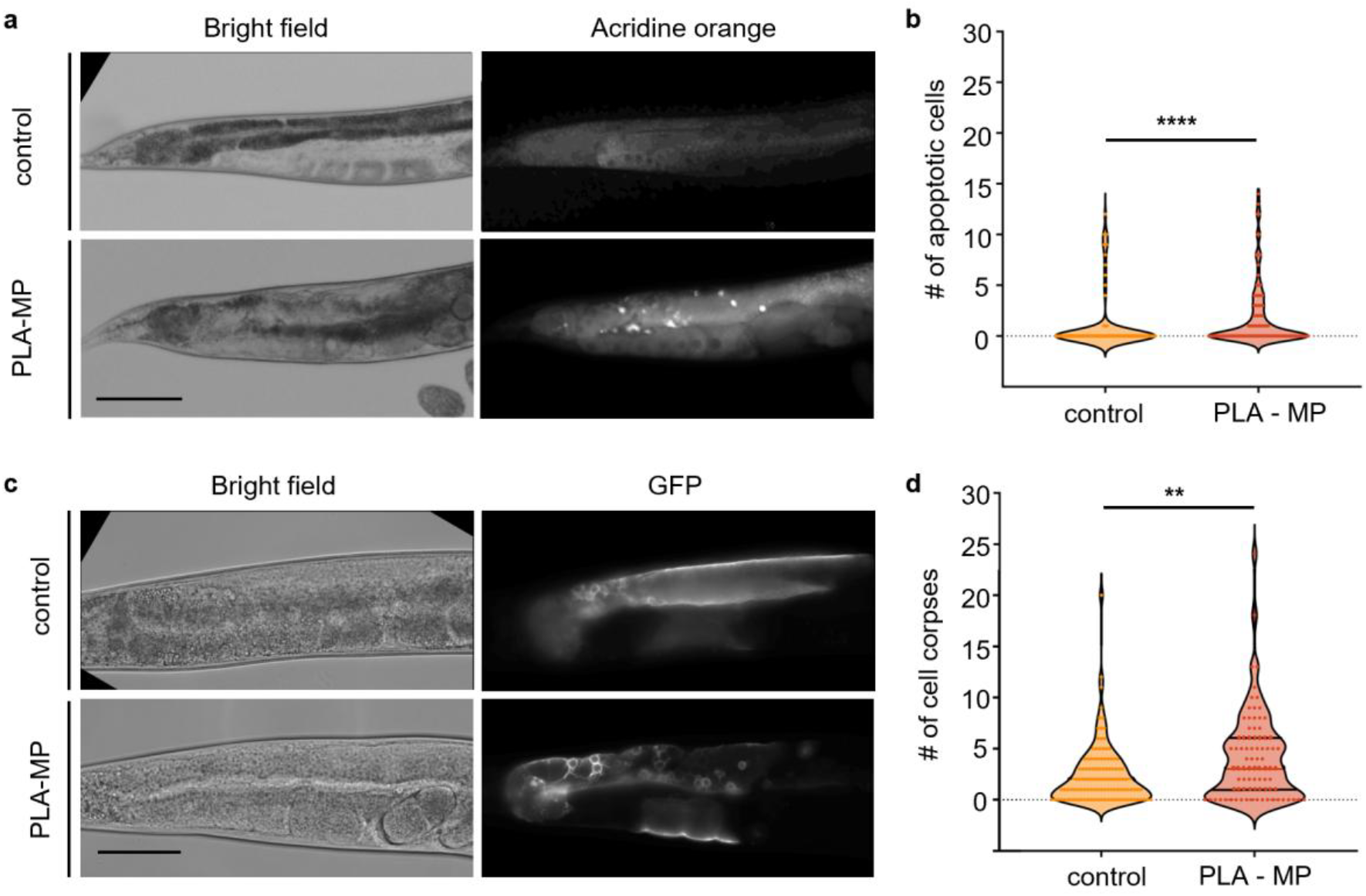
Increased apoptotic cells and cell death in the germline of PLA-exposed *C. elegans*. (**a**) Representative images of acridine orange staining in adult *C. elegans* fed *E. coli* with and without PLA-MPs. Scale bar = 100 µm. (**b**) Quantification of acridine orange staining of apoptotic cells in *C. elegans* with exposure to their regular E. coli diet only or with 1 µg/L PLA-MPs. N = 3, n = 102, 107. (**c**) Representative images of MD701, ced-1::GFP expression in adult *C. elegans* fed *E. coli* with and without PLA-MPs. Scale bar = 50 µm. (**d**) Quantification of MD701, ced-1::GFP apoptotic cells in *C. elegans* with exposure to their regular *E. coli* diet only or with 1 µg/L PLA-MPs. N = 4, n = 115, 84. Mann-Whitney test was used for statistical analysis. (** *p* ≤ 0.01; **** *p* ≤ 0.0001).

**Figure 6.**
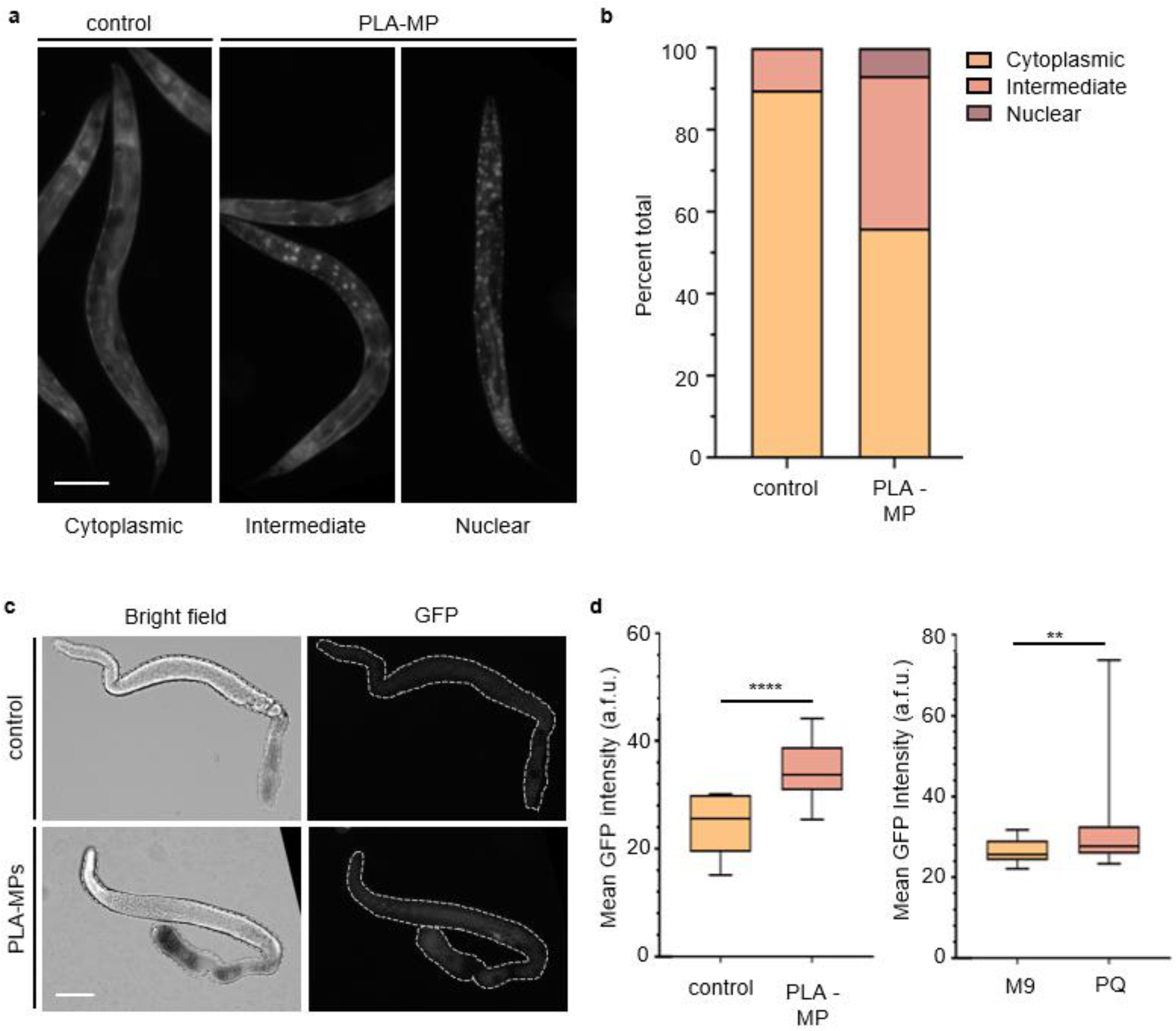
Multiple stress response pathways are activated in *C. elegans* after exposure to PLA-MPs. (**a**) Representative images of DAF-16::GFP expression in adult *C. elegans*, showing cytoplasmic (left), intermediary (center), and nuclear localization (right). Scale bar = 100 µm. (**b**) Quantification of DAF-16::GFP localization in adult *C. elegans* with exposure to their regular *E. coli* diet or with 1 µg/L PLA-MPs. N = 3, n = 122, 210. A chi-square test was used for statistical analysis (**** *p* < 0.0001). (**c**) Representative images of gst-4p::GFP in a dissected gonad from *C. elegans* without and with exposure to PLA-MP. The dotted line indicates the shape of the gonad. Scale bar = 50 µm. (**d**) Quantification of the mean GFP intensity from *C. elegans* expressing gst-4p::GFP without and with exposure to PLA-MP. a.f.u. = arbitrary fluorescence units. Error bars represent standard deviation (SD). **** *p* < 0.0001, ** *p* = 0.0100 by the two-tailed Mann-Whitney test, 95% C.I. n = 18–46 gonads per condition. N = 3, biological replicates.

### 3.4. PLA-MP Exposure Leads to DAF-16-Induced Stress in C. elegans and Oxidative Stress in the Gonad

PLA has been found to cause inflammation and increase reactive oxygen species (ROS) levels [24, 28, 29, 31], both of which are often associated with DNA damage [28], cell death [62], and reproductive decline [63, 64]. Additionally, a wide range of MP types, sizes, and exposure levels have been shown to cause stress via DAF-16/FOXO signaling and gst-4 [17, 23]. These pathways are central to the MP-induced stress response. Thus, we next asked whether the DAF-16 or the gst-4 stress responses were activated upon exposure to secondary PLA-MPs. Exposure of *C. elegans* to PLA-MPs increased the percentage of *C. elegans* with intermediate and nuclear expression of DAF-16 (*p* < 0.0001) (Figure 6a, b). We next asked whether oxidative stress was playing a role in the germline. We used an oxidative stress response reporter strain, *gst-4p::GFP*, in which GFP expression is driven by the glutathione S-transferase 4 (gst-4) promoter [65]. When this reporter strain was exposed to 1 µg/L PLA-MPs, we observed a significant increase in mean GFP signal intensity in the gonads (*p* < 0.0001) (Figure 6c, d). This response was similar to that observed with paraquat, an herbicide that induces oxidative stress (Figure 6d) [66, 67].

### 3.5. Exposure to Polylactic Acid Microplastics Does Not Reduce Lifespan

MP exposure shows varying effects on lifespan, which may be due to polymer type, size, and concentration. In many cases, MP exposure has been demonstrated to reduce lifespan in *C. elegans* [17, 23, 34, 68]. Additionally, activation of the DAF-16 stress response can be associated with increased lifespan [69-72]. Extended lifespan induced by stress events is linked to multiple pathways, including the insulin/insulin-like growth factor (IGF-1) receptor signaling pathway and diet [72]. Due to mixed reports on longevity in *C. elegans* and the potential for larger (> 5 µm) particles to obstruct the pharynx and thus alter nutrition [45], as well as studies indicating intestine-specific stress due to MP exposure [73], we next asked whether PLA-MPs would change the lifespan in *C. elegans*. Exposure of *C. elegans* to PLA-MPs did not alter their lifespan (Figure 7 and Supplemental Figure 3). There are only a limited number of studies on the lifespan of MPs composed of PLA or bio-based plastics. PLA-MPs reduced lifespan in the rotifer *Brachionus plicatilis*, and leachates from bioplastics reduced lifespan in *C. elegans* [29, 74]. These studies used different concentrations and types of PLA-MPs and their leachates [29]. In contrast to the *B. plicatilis study*, Shao et al. reported a reduction in lifespan with 3 µm-diameter PLA-MPs, but only at higher exposure levels (100 µg/L) [29]. The varying effects on lifespan may be due to the PLA-MP relative exposure, and properties of the particles.

**Figure 7.**
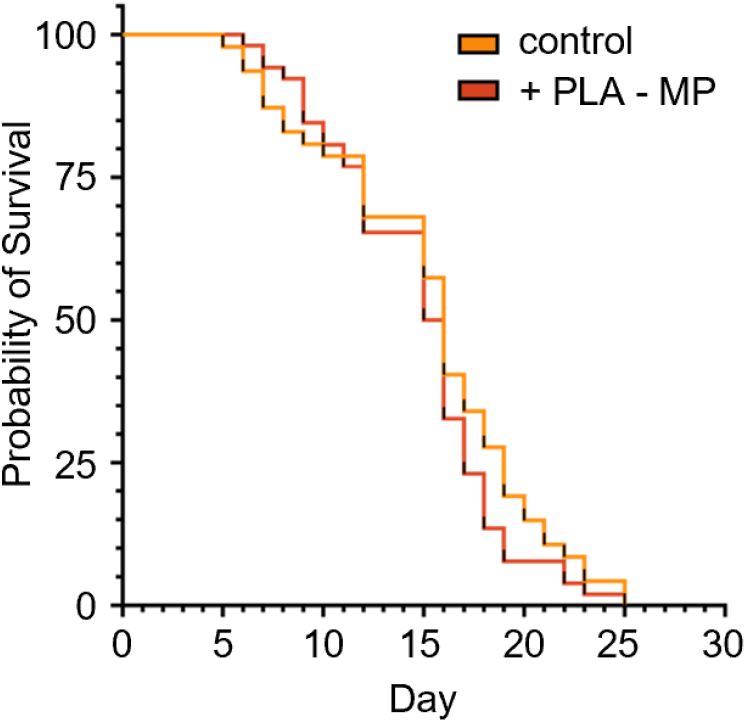
Lifespan of *C. elegans* exposed to PLA-MPs. Survival assay for *C. elegans* exposed to *E. coli* (no MPs) or 1 µg/L PLA-MPs continuously. n = 47, 52. Not significant, p = 0.249. Log-rank (Mantel–Cox).

## 4. Conclusions

Here we show that PLA-MPs, like soluble toxicants, can trigger germline damage and apoptosis. Therefore, irregular secondary PLA-MPs are emerging pollutants that can affect fertility and germline health. Our work complements the emerging literature on pollutants with similar endpoints for particle-based environmental pollutants rather than soluble chemicals. We presented a method to create and mimic environmentally relevant PLA-MPs using a cryo-mill method to examine physiological defects and their underlying cause. Exposure of *C. elegans* to a heterogeneous pool of 1 µg/L PLA-MPs (varying in size, shape, and surface texture) reduces fertility but not lifespan. Exposure by ingestion is potentially as low as 30 ng/L, demonstrating defects in fertility at far below commonly accepted levels of MP exposure. While we did not observe an effect on lifespan from PLA-MP exposure, we did observe a drastic reduction in fertility and changes in the germline. The germline is likely more sensitive than somatic tissues under our exposure conditions. The reduction in fertility is driven by alterations to the germline, characterized by chromosome aberrations, oxidative stress, and cell death. We focused on using mechanically processed secondary MPs to better mimic those found in the environment. In reality, human and animal MP exposure is represented by various polymer types, mixtures (polymer and additive), absorption of environmental chemicals, and multiple methods of degradation (mechanical, UV, chemical, etc.). Future work can methodically add UV aging and chemical exposure to simulate more advanced environmental degradation.

## Supplementary Materials

Supplemental Figure 1: Frequency distribution of microplastic sizes.

Supplemental Figure 2: Number of eggs hatched, unhatched, and number of males produced per brood in *C. elegans* exposed to PLA MPs.

Supplemental Figure 3: Lifespan of *C. elegans* exposed to PLA-MPs.

## Author Contributions

Conceptualization, J.C.H.; methodology, J.C.H., C.A.O.M., D.M.M., R.R., and R.L.P.; validation, J.C.H. and R.L.P.; formal analysis, J.C.H., C.A.O.M., and R.L.P.; investigation, J.C.H., C.A.O.M., D.F., R.R., D.M.M., M.F.G., L.R.S., V.P., A.G., and R.L.P.; resources, J.C.H.; data curation, J.C.H., C.A.O.M. and D.M.M.; writing—original draft preparation, J.C.H. and C.A.O.M.; writing—review and editing, J.C.H.; visualization, J.C.H. and C.A.O.M.; supervision, J.C.H.; project administration, J.C.H.; funding acquisition, J.C.H. All authors have read and agreed to the published version of the manuscript.

## Data Availability Statement

The raw data supporting the conclusions of this article will be made available by the authors on request.

## Acknowledgments

Many undergraduate student researchers have helped move this project forward, and we thank them for their time and valuable input. Thank you to Dr. Oxley for her advice and discussion. We thank C. James for helping collect data on brood size during her training in the lab. We also thank St. Mary’s University and Texas State University for their support. St. Mary’s University funds from the Department of Biological Sciences, the Benjamin F. Biaggini Endowment, and the Office of Student Research and Inquiry SURF program. *C. elegans* strains were provided by the CGC, which is funded by the NIH Office of Research Infrastructure Programs (P40 OD010440).

## Abbreviations

*C. elegans*: *Caenorhabditis elegans*
DAF-16: Forkhead box O (FOXO) protein
*E. coli*: *Escherichia coli*
GFP: Green fluorescent protein
GST-4: Glutathione S-transferase 4
LA: Lactic acid
L1: Larval stage 1
L4: Larval stage 4
MP: Microplastic
NGM: Nematode Growth Media Agarose
NP: Nanoplastic
PBS: phosphate-buffered saline
PFA: paraformaldehyde
PLA: Polylactic acid
PS: Polystyrene
PQ: Paraquat
ROS: Reactive oxygen species
SEM: Scanning Electron Microscopy
UV: Ultraviolet

