## Supplementary material for "Environmentally Relevant Polylactic Acid Microplastics from 3D Printing Induce Germline Apoptosis and Reproductive Decline in *Caenorhabditis elegans*": Supplimental Figures

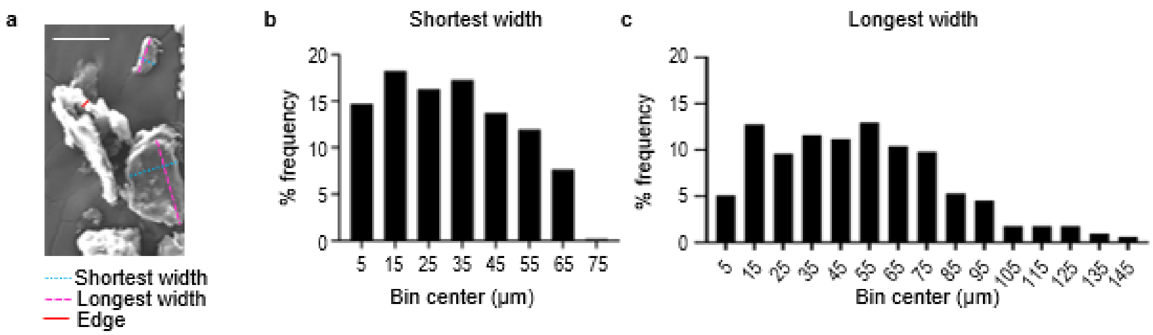


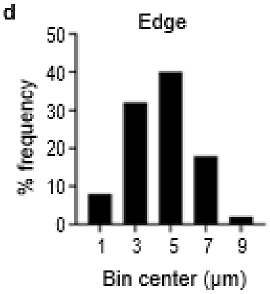


e

| | Shortest width (um) | Longest width (um) | Edge (um) |
|---|---|---|---|
| Total number of values | 510 | 510 | 50 |
| Minimum | 1.90 | 2.25 | 1.14 |
| 25% Percentile | 15.93 | 26.76 | 2.90 |
| Median | 30.22 | 49.613 | 5.07 |
| 75% Percentile | 46.74 | 72.05 | 5.89 |
| Maximum | 71.92 | 146.37 | 8.50 |
| Mean | 31.50 | 51.80 | 4.59 |
| Std. Deviation | 18.11 | 30.52 | 1.75 |
| Std. Error of Mean | 0.80 | 1.35 | 0.24 |
| Lower 95% CI of mean | 29.92 | 49.15 | 4.09 |
| Upper 95% CI of mean | 33.08 | 54.46 | 5.09 |

**Supplemental Figure 1:** Frequency distribution of microplastic sizes. (**a**) Representative SEM image of PLA-MPs and diagram showing how MPs were measured. Scale bar = 50 µm. (**b**) Frequency distribution of PLA-MPs measured in the smallest direction, distributed into µm bins. n = 510. (**c**) Frequency distribution of PLA-MPs measured at their longest point, distributed into 10 µm bins. n = 510. (**d**) Frequency distribution of PLA-MPs measured at their edge, distributed into 2 µm bins. n = 50. (**e**) Descriptive statistics for PLA-MP size measurements.

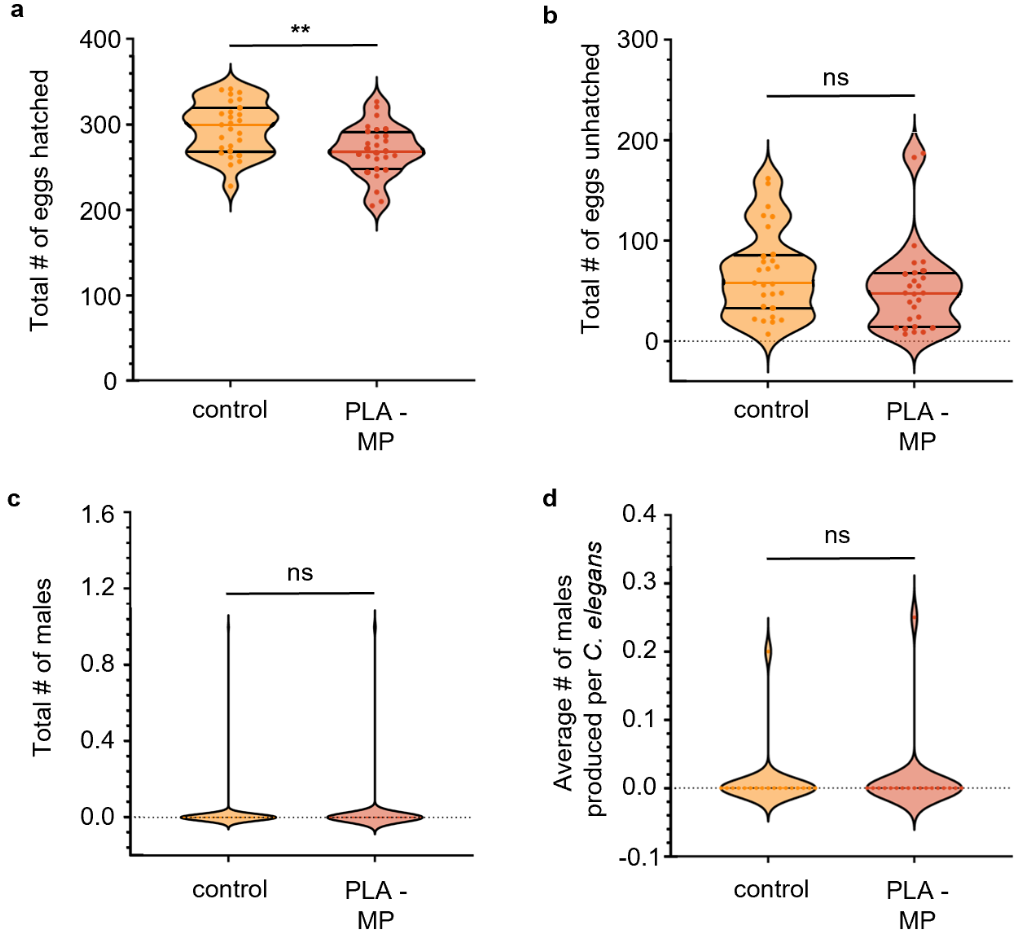


**Supplemental Figure 2**: Number of eggs hatched, unhatched, and number of males produced per brood in *C. elegans* exposed to PLA-MPs. (**a**) Number of eggs hatched, N = 4, n = 27, 27. (**b**) Number of eggs unhatched, N = 4, n = 27, 27. (**c**) Number of males N = 2, n = 126, 126. (**d**) Average number of males produced from parental *C. elegans*. N = 2, n = 18, 18. Mann-Whitney (two-sample) tests were used for statistical analysis (** $p \leq 0.01$).

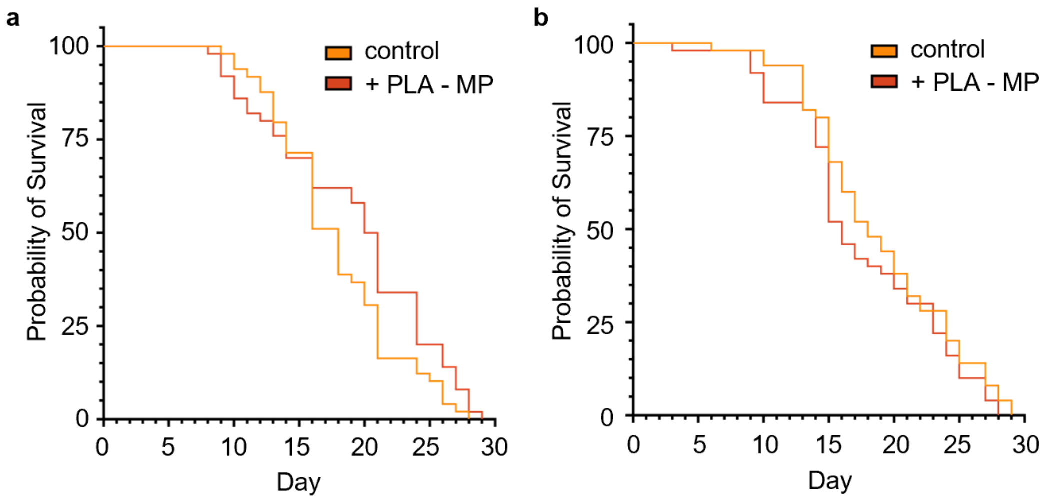


**Supplemental Figure 3:** Lifespan of *C. elegans* exposed to PLA-MPs. (**a**) Replica of survival assay for *C. elegans* exposed to *E. coli* (no MPs) or 1 µg/L PLA MPs continuously. n = 49, 50. Not significant, $p$ = 0.059. Log-rank (Mantel–Cox). (**b**) Replica of survival assay for *C. elegans* exposed to *E. coli* (no MPs) or 1 µg/L PLA MPs continuously. n = 50, 50. Not significant, $p$ = 0.259. Log-rank (Mantel–Cox).
